# The dynamics of nucleolus-centromeres interaction in living cells

**DOI:** 10.64898/2026.04.13.718322

**Authors:** Wei-Hong Yeo, Emma Freeman, Alex B. Willis, Hao F. Zhang, Daniel Foltz, Sui Huang

**Affiliations:** Department of Biomedical Engineering, Northwestern University, Evanston, IL, USA; Department of Cell and Developmental Biology, Northwestern University, Chicago, IL, USA; Department of Biochemistry and Molecular Genetics, Northwestern University, Chicago, IL, USA

**Keywords:** nucleoli, centromeres, live cell imaging, 4D dynamics

## Abstract

Nucleoli and centromeres play essential roles in cellular proliferation and homeostasis, and are structurally and functionally interconnected. Centromeres frequently cluster around nucleoli, and some centromere assembly factors are known to reside in the nucleoli. To investigate the spatial and temporal relationships between these nuclear domains, we examined their dynamics in living cells. We imaged HeLa cells stably expressing mCherry-NPM1 and GFP-CENP-A using time-lapse microscopy. The results show that a subset of centromeres exhibits dynamic behavior during interphase, migrating over micrometer-scale distances within two hours. On average, 40–50% of centromeres maintain an association with nucleoli throughout interphase, with some cells displaying nucleolar-centromere association and dissociation within hours. Upon entry into mitosis, nucleoli are disassembled, and NPM1 localizes to the periphery of mitotic chromosomes. Nucleolar-centromere interactions are re-established in early G1, coinciding with the assembly of new centromeres. Treatment with actinomycin D, an inhibitor of RNA polymerase I, significantly reduces nucleolar size, nucleolar-centromere interactions, and centromere dynamics. Furthermore, post-mitotic nucleolar reformation is impaired. These findings highlight the dynamic nature of centromeres in interphase nuclei and their interactions with nucleoli. This behavior is partially dependent on rDNA transcription and nucleolar integrity, underscoring the critical roles of nucleoli, centromeres, and their interaction in 4D genome organization.

## INTRODUCTION

Nucleoli and centromeres are essential nuclear structures that play critical roles in cell proliferation and maintaining genomic integrity. Centromeres are distinct chromatin domains that orchestrate chromosome segregation during mitosis. Centromere protein A (CENP-A) replaces histone H3 to form a stable centromeric nucleosome found primarily at the centromere (Palmer et al., 1991; Sullivan et al., 1994; Warburton et al., 1997; Meluh et al., 1998; Yoda et al., 2000; Régnier et al., 2005; Foltz et al., 2006; Liu et al., 2006). Human CENP-A directs the recruitment of the constitutive centromere-associated network (CCAN) comprised of at least 17 CENPs (Foltz et al., 2006; Okada et al., 2006; Hori et al., 2008). The composition of this macromolecular complex expands greatly as cells move from interphase into mitosis, when the complex grows to over 80 proteins involved in regulating and sensing microtubule attachment and facilitating chromosome movement during cell division (Musacchio and Salmon, 2007; Verdaasdonk and Bloom, 2011; McKinley and Cheeseman, 2016). The location of the centromere in higher eukaryotes is determined epigenetically, independently of DNA sequence (Cleveland et al., 2003; Allshire and Karpen, 2008; Black and Bassett, 2008). The formation of neocentromeres at non-centromeric loci in humans, and the activation of neocentromeres in experimental systems demonstrate that DNA sequence is not required for centromere activity (Cook et al., 1997; Ishii et al., 2008; Marshall et al., 2008; Hori et al., 2012). CENP-A forms a stable nucleosome at the centromere; therefore, it is widely accepted to be the epigenetic identifier of the centromere (Heun et al., 2006; Barnhart et al., 2011; Guse et al., 2011; Mendiburo et al., 2011). While it is well known that centromeres are the key engines for chromosome segregation during mitosis, the role of centromeres in interphase cells is much less understood.

Nucleoli serve as the primary site of ribosome synthesis, facilitating and regulating this complex, multi-step process (Busch and Smetana, 1970; Pederson, 1998, 2011; Dörner et al., 2023). Ribosome biogenesis involves transcription by RNA polymerase I (Pol I) using rDNA as the template to produce polycistronic pre-rRNAs. Pre-rRNAs contain external and intrinsic sequences that are removed to generate 18S, 5.8S, and 28S rRNAs in human cells. The processed rRNAs are assembled with corresponding large and small subunit ribosomal proteins to form 40S and 60S pre-ribosomal particles, which are subsequently exported to the cytoplasm where they undergo further maturation processes before becoming fully functional ribosomes for protein translation (Hadjiolov, 1985; Leary and Huang, 2001; Pelletier et al., 2018; Shore and Albert, 2022). Nucleoli are also multi-functional organelles involved in cellular processes that extend beyond ribosome synthesis (Pederson, 1998; Boisvert et al., 2007; Pederson, 2011; Iarovaia et al., 2019). Many cellular activities are associated with nucleoli, including E3 ubiquitin-protein ligase MDM2 mediated regulation of p53 (Deisenroth and Zhang, 2011; Liu et al., 2016), stress response (Boulon et al., 2010; James et al., 2014), macromolecular trafficking, assembly of various ribonucleoprotein particles, microRNA metabolism (Politz et al., 2009), genome organization and others (Dubois and Boisvert, 2016; Pederson, 1998; Németh et al., 2010; van Koningsbruggen et al., 2010; Guetg and Santoro, 2012; Bersaglieri and Santoro, 2019; Vertii et al., 2019; Bersaglieri et al., 2022).

While centromeres and nucleoli have been extensively studied individually for their roles in cell proliferation and function, an integral relationship exists between the two subnuclear structures across species. Centromeres cluster around nucleoli in many cell types. The association of centromeres with the nucleolus is dynamic and related to the transcriptional status of the centromeric DNA (Bury et al., 2020; Rodrigues et al., 2023). Undifferentiated cell types and cancer cells show higher degrees of association between centromeres and nucleoli (Rodrigues et al., 2023). Factors, including Holliday Junction-Recognition Protein (HJURP) and Nucleophosmin 1 protein (NPM1), involved in centromere assembly are enriched in centromeres and nucleoli (Dunleavy et al., 2009; Foltz et al., 2009). In Drosophila cells, the position of the centromere around the nucleoli is driven by the nucleophosmin family protein nucleoplasmin-like protein (NLP) and Modulo (Padeken et al., 2013). These observations suggest a functional connection between these two nuclear structures in interphase cells. To further explore this relationship, we examined the temporal and spatial dynamics of centromeres and nucleoli, as well as their interactions, in living cells using time-lapse microscopy.

## MATERIAL AND METHODS

### Cell Culture

HeLa cells were infected with lenti-viral expression constructs containing fusion proteins of mCherry-NPM1 and eGFP-CENP-A (Fig. 1A). The cells stably expressing these proteins were evaluated for their physiological behaviors and they showed normal growth conditions. Cells were grown in DMEM supplemented with 10% FBS and antibiotics. For imaging, cells were plated in glass bottom 30-mm dishes and placed on an inverted confocal microscope (Nikon; A1+) with complete environmental controls (CO_2_ and heating). For the actinomycin D (ActD) treatment, 0.04 µg/mL, drug was added to the culture media immediately before being placed onto the microscope and cells were selected and imaged over time.

**Figure 1.**
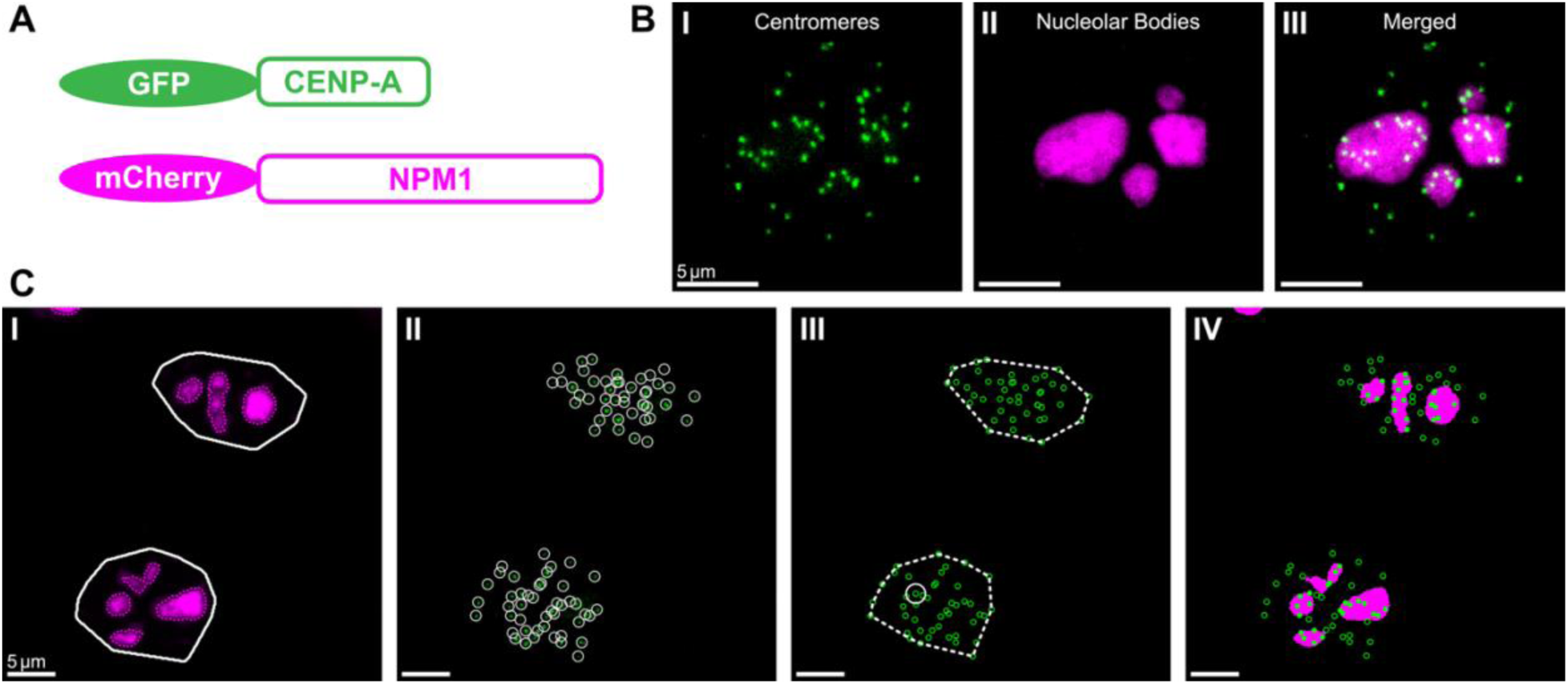
Construction of reporter cell lines and image processing workflow. **A.** Stick diagram of the construction of mCherry-NPM1 and GFP-CENP-A, lenti viral infection and selections. **B.** Interphase cells showing **I.** centromeres, **II.** NPM1, and **III.** both channels merged. **C.** Illustration of image processing steps. **I.** Fractional area occupied by nucleolar bodies, **II.** identification of centromeres, **III.** centromere-centromere clustering using Ripley’s L, and **IV.** association of centromeres with the nucleolus.

### Image Acquisition

Time-lapse fluorescence microscopy of centromere- and nucleolus-stained cells was performed using a confocal microscope, equipped with a 60x objective (Nikon; Plan Apo λ 60x Oil) and a GaAsP detector. The imaging setup was controlled via Nikon NIS-Elements software. Cells were maintained at 37°C with 5% CO_2_ in a live-cell incubation chamber to ensure optimal physiological conditions throughout the imaging period.

HeLa cells were cultured in glass-bottom 30 mm dishes and imaged at intervals of 7.5 or 15 minutes over a total duration of 20 hours, using a seven-section *z*-stack with autofocus activated at each time point. Seven z-sections for each image sample were obtained under a perfect focus mode to capture most of the centromeres from each cell. Samples were illuminated with laser excitation wavelengths of 488 nm and 561 nm to detect GFP-and mCherry-labeled structures, respectively, with emission signals collected using optimized bandpass filters (Fig. 1B). Images were captured at time intervals of 800 or 1200 seconds, depending on the experimental setup.

For subsequent analysis, *z*-stacks were flattened using maximal projection to generate 2D images, ensuring that most of the centromeres in each cell were represented. Since nucleoli are the thickest nuclear structures in the *z*-dimension, centromeres appearing above or below the nucleoli are likely near nucleoli bodies, as there is minimal free nuclear space between nucleoli and the nuclear envelope (Fig. 1B).

### Image Processing

From the video files, cells undergoing mitosis were manually identified based on morphological criteria and fluorescence intensity changes. Selected cells were cropped to isolate relevant regions and time points, generating sub-images for further processing. Image cropping was performed manually, ensuring that all selected regions retained sufficient spatial resolution and fluorescence signal for subsequent analysis.

The image processing workflow was implemented in MATLAB 2023b, where raw videos corresponding to the nucleolar and centromere markers were initially loaded and pre-processed. To enhance image quality, intensity normalization was applied through histogram stretching, ensuring that pixel intensity values were scaled within the 10^th^ and 99^th^ percentiles of the original distribution. To reduce noise and improve segmentation, Gaussian filtering was employed with a smoothing parameter of 1 for nucleolar images and 3 for centromere images. Additional noise reduction in the centromere images was achieved using median filtering with a kernel size of 3×3 pixels.

Binary segmentation was performed using adaptive thresholding to identify nucleolar, nuclear, and centromere regions. The nucleolar signal was binarized using a manually determined threshold, while nuclear regions were extracted by applying a Gaussian smoothing operation followed by thresholding at a manually determined threshold. Similarly, centromere regions were identified using a combination of median filtering and thresholding at a cutoff value manually determined. Post-processing included morphological operations such as dilation with a circular structuring element of radius 5 pixels, and closing with a structuring element of radius 10 pixels to fill gaps in the nuclear and centromere masks. Small objects with an area of fewer than 5000 pixels were removed to eliminate false positives. The boundaries of nuclear and centromere regions were further refined using convex hull operations to ensure accurate area computations. (Fig. 1CI)

After segmentation, objects of interest were detected and quantified. Nucleolar structures were identified by computing connected components within the binarized nucleolar mask, and their centroids and areas were extracted. To obtain better estimates of centromere positions, we used the ThunderSTORM plug-in (Ovesný et al., 2014) on ImageJ to identify coordinates of the centromeres in the images (Fig. 1CII), which were then exported as a .csv file for further processing in MATLAB. To classify centromeres as being within the nucleolus or the nuclear space, an 8-connected neighborhood search was performed, allowing the detection of centromeres that were in direct contact with the nucleolus.

A range of morphological and spatial metrics were computed for each cell at each time point. The convex hull area of centromeres was determined by identifying the smallest convex polygon enclosing all centromeres within a given nucleus, and the enclosed area was computed accordingly (Fig. 1CIII). The clustering of centromeres was quantified using Ripley’s *L*-function (Ripley, 1976) with a radius parameter of 10 pixels, with values adjusted to remove expected clustering effects under complete spatial randomness (Fig. 1CIII, Supp. Note 1, Supp. Fig. 1). The same approach was applied to nucleolar bodies to determine their convex hull area. The fraction of centromeres residing within the nucleolus was calculated as the proportion of centromeres classified as nucleolus-associated relative to the total centromere count in the nucleus (Fig. 1CIV). The total nucleolar area was determined by summing the areas of individual nucleolar bodies detected within the binarized nucleolar mask.

### Statistical Analyses

To quantitatively assess nuclear, nucleolar, and centromeric structures, we measured several morphological and spatial features. The degree of centromere clustering was measured using Ripley’s *L*-function, providing an estimate of how centromeres deviated from a random spatial distribution. The fraction of centromeres within the nucleolus was calculated as the ratio of nucleolus-associated centromeres to the total number of centromeres within a given cell. Using the microscope’s calibrated pixel size, the total nuclear area was extracted from the binarized nuclear mask, while the centromere convex hull area was computed based on the convex polygon enclosing all centromeres in the nucleus. Similarly, the nucleolar convex hull area was determined using the convex polygon enclosing nucleolar bodies.

The absolute nucleolar area was calculated as the sum of the areas of individual nucleolar bodies within the nucleus. The total number of centromeres was counted from the .csv dataset, while the number of nucleolar bodies was obtained from the segmented nucleolar mask. Centromere density was defined as the number of centromeres per unit nuclear area, and nucleolar body density was computed as the number of nucleolar bodies per unit nuclear area. To assess the spatial organization of centromeres and nucleolar bodies within the nucleus, the fractional nuclear area occupied by centromeres and nucleolar bodies was computed by normalizing their respective convex hull areas by the total nuclear area. Additionally, the absolute nucleolar area fraction was calculated as the ratio of the total nucleolar area to the nuclear area.

To evaluate centromere positioning relative to nuclear and nucleolar boundaries, the mean distances of centromeres to the nucleolus and nucleus were computed. Distance transform algorithms were applied to both the nuclear and nucleolar masks to generate distance maps, which provided the Euclidean distance of each pixel from the nearest nucleolar or nuclear boundary. To improve spatial resolution, these masks were upscaled by a factor of 10, and the centroid coordinates of centromeres were mapped onto the transformed space to extract their precise distances from the nucleolus and nucleus.

To analyze temporal trends in centromere and nucleolar dynamics, data from individual time-lapse experiments were aggregated and smoothed to facilitate comparisons between experimental conditions. The dataset included multiple experimental files, each containing centroid trajectories across time points. Data selection was performed based on the experimental metadata, where only files marked for processing were included.

For each dataset, extracted morphological and spatial features were interpolated onto a common time grid ranging from 6 to 12 hours, with an interpolation interval of 400 seconds. This temporal alignment ensured that all trajectories were analyzed on a uniform time axis, facilitating direct comparisons between experimental conditions. Interpolation was performed using a modified Akima cubic interpolation method, which preserves local shape characteristics while avoiding oscillatory artifacts in sparsely sampled data.

To reduce noise while preserving biologically relevant trends, the Whittaker-Eilers smoother was applied to the interpolated data. This approach balances smoothness and fidelity to the original data by solving a penalized least squares problem, where a smoothing parameter (*λ* = 10) controls the tradeoff between fitting the data and imposing smoothness constraints. Additionally, a second-order difference penalty (Δ = 2) was used to regulate the smoothness of the fitted curve while preventing over-smoothing of transient biological variations. Prior to smoothing, missing data points were identified and excluded from the fitting process, ensuring that only valid observations contributed to the final smoothed trajectories.

To assess dynamic changes in centromere and nucleolar features over time, individual track trajectories were plotted alongside condition-averaged trends. A set of time-course plots was generated for each extracted feature, displaying both raw interpolated trajectories and Whittaker-Eilers smoothed curves. The time axis was centered such that *t* = 0 corresponded to the beginning of anaphase, enabling alignment of trajectories relative to this reference point.

## RESULTS

### 4D Centromeres dynamics in the interphase cell nucleus

The factors that influence the dynamic localization of centromeres within the nucleus is not well understood. To visualize centromeres, nucleoli, and their association in living cells, we constructed a HeLa cell line stably expressing eGFP-CENP-A and mCherry-NPM1 (Fig. 1A). Generally, the cell line propagates well, and nucleoli (mCherry) and CENP-A (GFP) signals are robust, and image acquisition can be achieved within milliseconds using Nikon confocal A1 microscope (Fig. 1B). To quantify centromere and nucleolar organization, we segmented nucleolar, nuclear, and centromere regions using adaptive and manual thresholding methods (Fig. 1CI). Morphological operations and convex hull refinement were applied to improve mask accuracy. Small artifacts were excluded based on area thresholds. Nucleolar structures were quantified by connected component analysis, and centromeres were localized using ThunderSTORM (Fig. 1CII), followed by classification as nucleolar- or nuclear-associated using neighborhood-based detection. We computed various morphological and spatial metrics, including convex hull areas and Ripley’s *L*-function to assess clustering (Fig. 1CIII, Supp. Note 1, Supp. Fig. 1). The proportion of nucleolar-associated centromeres and total nucleolar area were also measured (Fig. 1CIV).

Cells were imaged up to 20 hours with 7.5 or 30 minutes intervals. As shown in Figs. 2AI to AVI, many centromeres move continuously during interphase, with displacements reaching several micrometers over the course of an hour (Fig. 2AII, 2AV; enlarged views in Fig. 2AIII, 2AVI; individual frames, single-particle tracks and time-lapse panels in Supp. Fig. 2). Centromeres were observed both moving toward and away from each other in individual cells, demonstrating their dynamic nature. The observed movement is unlikely to have resulted from global nuclear rotation, as centromere trajectories were not uniformly directed in one orientation.

**Figure 2.**
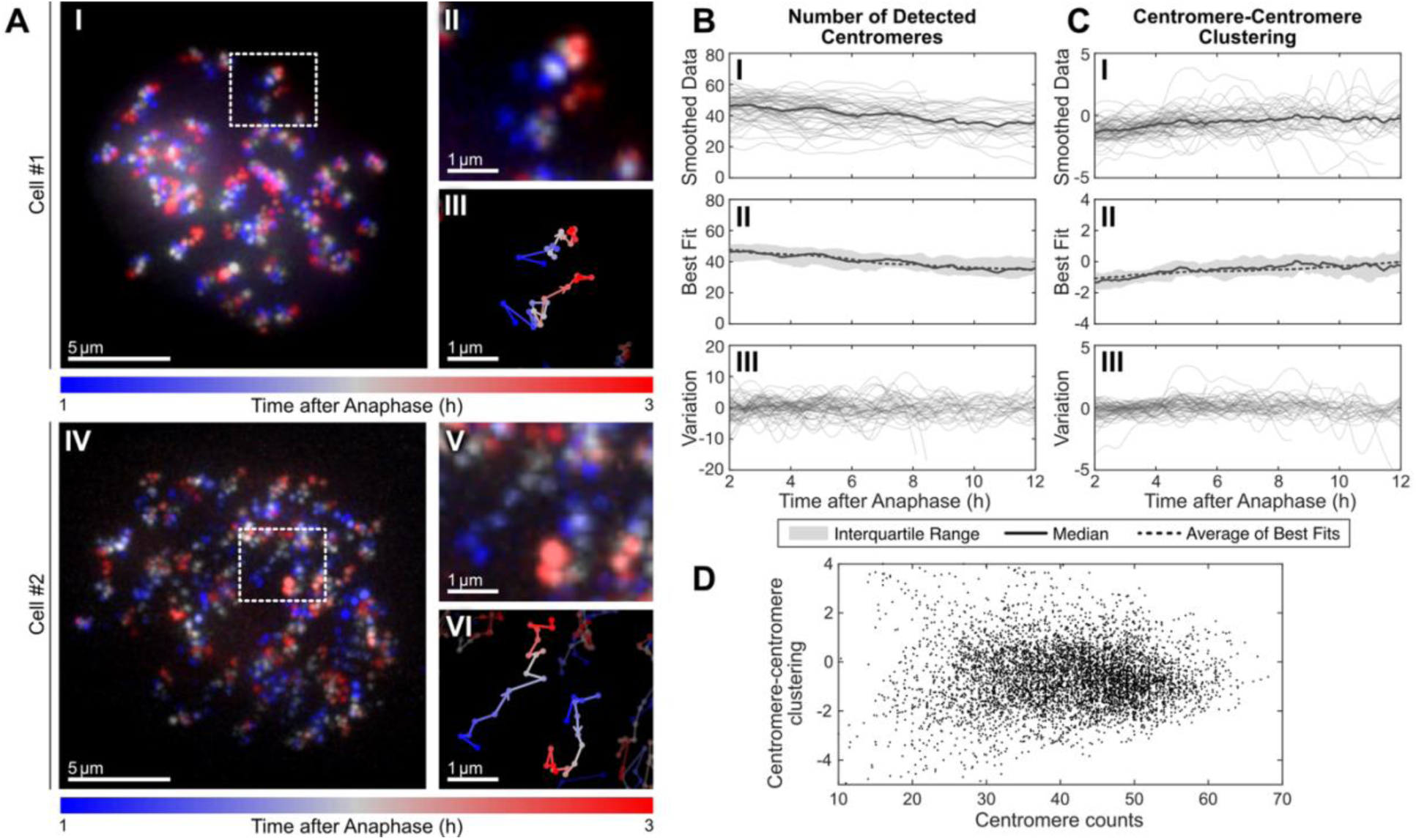
Dynamics of centromere distribution and clustering over time. **A.** Aligned centromeres distribution over time. **I, IV.** Color-coded time stack of centromeres of a cell over 2 hours, **II, V.** close-up view of 2 centromeres exhibiting long-range movement, **III, VI.** single-particle track of the 2 centromeres showing movement of up to 5 μm. **B.** Plots of number of detected centromeres, and **C.** centromere-centromere clustering, each illustrating **I.** smoothed raw data, **II.** best fit illustrating the trend, and **III.** the variation obtained from removing trend from the smoothed raw data. **D.** Plot of centromere-centromere clustering metrics against centromere counts.

To determine whether centromere dynamics are influenced by detected centromere number, we quantified centromere counts from live-cell images and applied smoothing techniques (detailed in Methods and Materials) to estimate their temporal changes for each cell (Fig. 2BI). Using best-fit analysis, we decomposed the data into trend (Fig. 2BII) and residual components (Fig. 2BIII), where the trend reflects overall centromere number changes, while residuals capture fluctuations over time. The trend analysis revealed a gradual decline in median centromere number from 48 at 2 hours to 36 at 12 hours post-anaphase (p < 0.001, denoted as ***, Supp. Fig. 4A), suggesting a slight reduction over time, which is most likely due to the loss of fluorescence from phototoxicity. It could also be due to centromere fusions. To test this idea, we analyzed dynamics using Ripley’s *L* function (Ripley, 1976) at a scale of 1 µm (Supp. Note 1, Fig. 2CI). Similarly, we used best-fit analysis to decompose the data into its trend (Fig. 2CII) and residual components (Fig. 2CIII). The trend demonstrates a progressive increase in centromere clustering, with Ripley’s *L* values rising from approximately −1 at 2 hours post-anaphase to 0 at 12 hours post-anaphase (*p* < 0.001, denoted as ***, Supp. Fig. 4B). These results indicate a progressive spatial reorganization of centromeres throughout interphase, with a tendency for them to localize close to one another.

### Centromeres are clustered around nucleoli

Previous studies, including those from our labs and others, have demonstrated that centromeres cluster around nucleoli, with the extent of clustering influenced by both physiological and pathological conditions (Bury et al., 2020; Rodrigues et al., 2023). However, whether the nucleolar-centromere association is dynamic in interphase cells over time is not evaluated. Here, we performed live-cell imaging to analyze centromere movement relative to nucleoli.

As shown in Fig. 3A, centromeres (green) exhibit clustering around nucleoli, and the association is not static. Live-cell imaging revealed that although a subset of centromeres exhibits dynamic behavior, periodically approaching and moving away from nucleolar bodies (Fig. 3A, arrows), the majority remains stably associated with nucleoli. This is further illustrated in Supp. Fig. 5III, where most centromeres are consistently either associated or unassociated with the nucleolus, with only a small subset spending intermittent periods near the nucleolus. We found that there are no significant differences in the rate of movement, variability of movement, and directional persistence for these centromeres relative to association with nucleolar bodies (Supp. Fig. 5). To further illustrate these dynamics, we also show a larger panel of time-lapse images (Supp. Fig. 6) which highlights the to-and-fro movement of centromeres relative to nucleoli.

**Figure 3.**
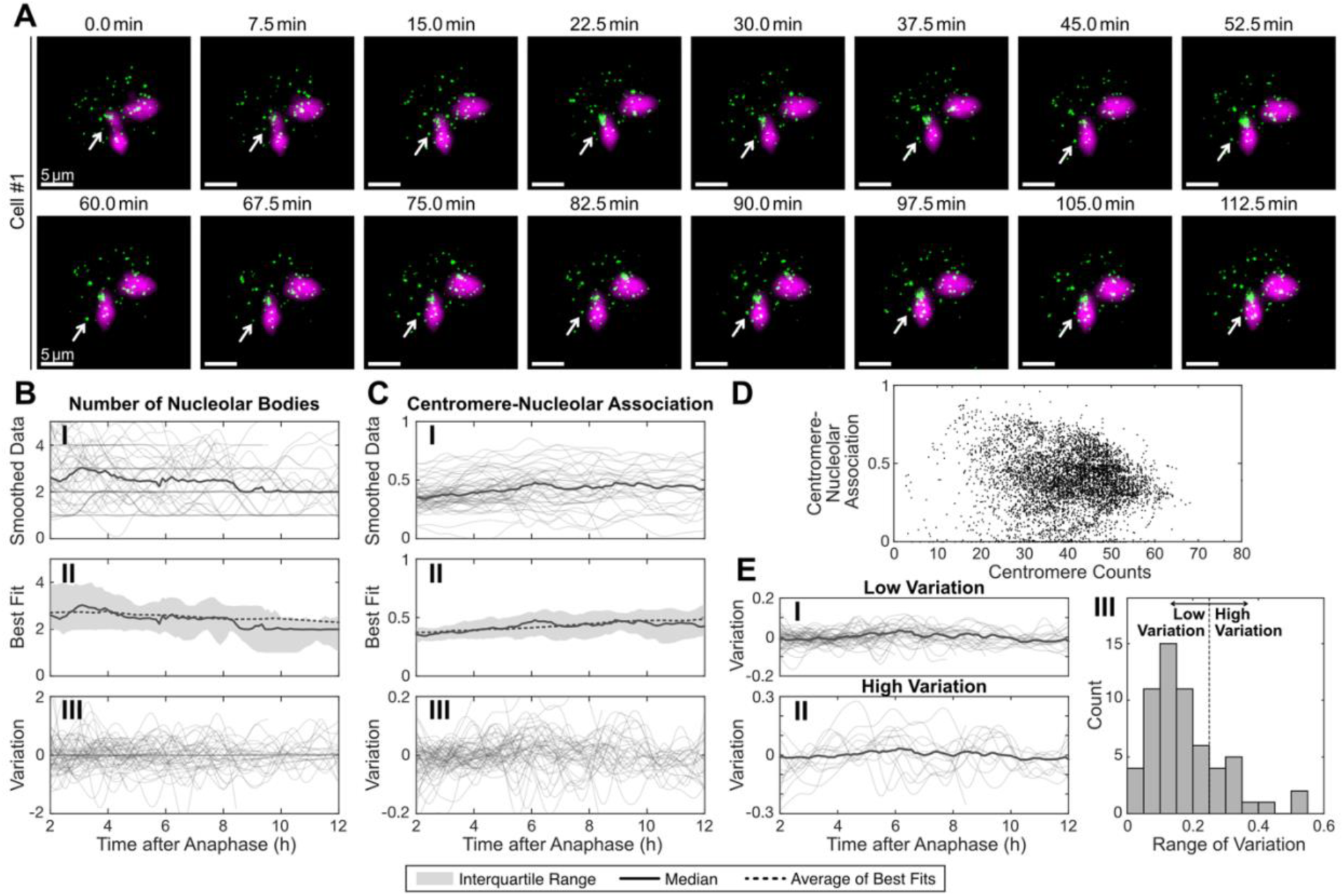
Centromere-nucleolus interactions and their variability. **A.** Panel of images illustrating the movement of centromeres relative to NPM1 over 2 hours. **B.** Plots of number of nucleolar bodies and **C.** centromere-nucleolar clustering, each illustrating **I.** smoothed raw data, **II.** best fit illustrating the trend, and **III.** the variation obtained from removing trend from the smoothed raw data. **D.** Plot of centromere-nucleolar association against centromere counts. **E.** Plot of the variation of centromere-nucleolar association for samples with **I.** low variations and **II.** high variations, and **III.** a histogram plot illustrating the number of samples with low and high range of variation.

**Figure 4.**
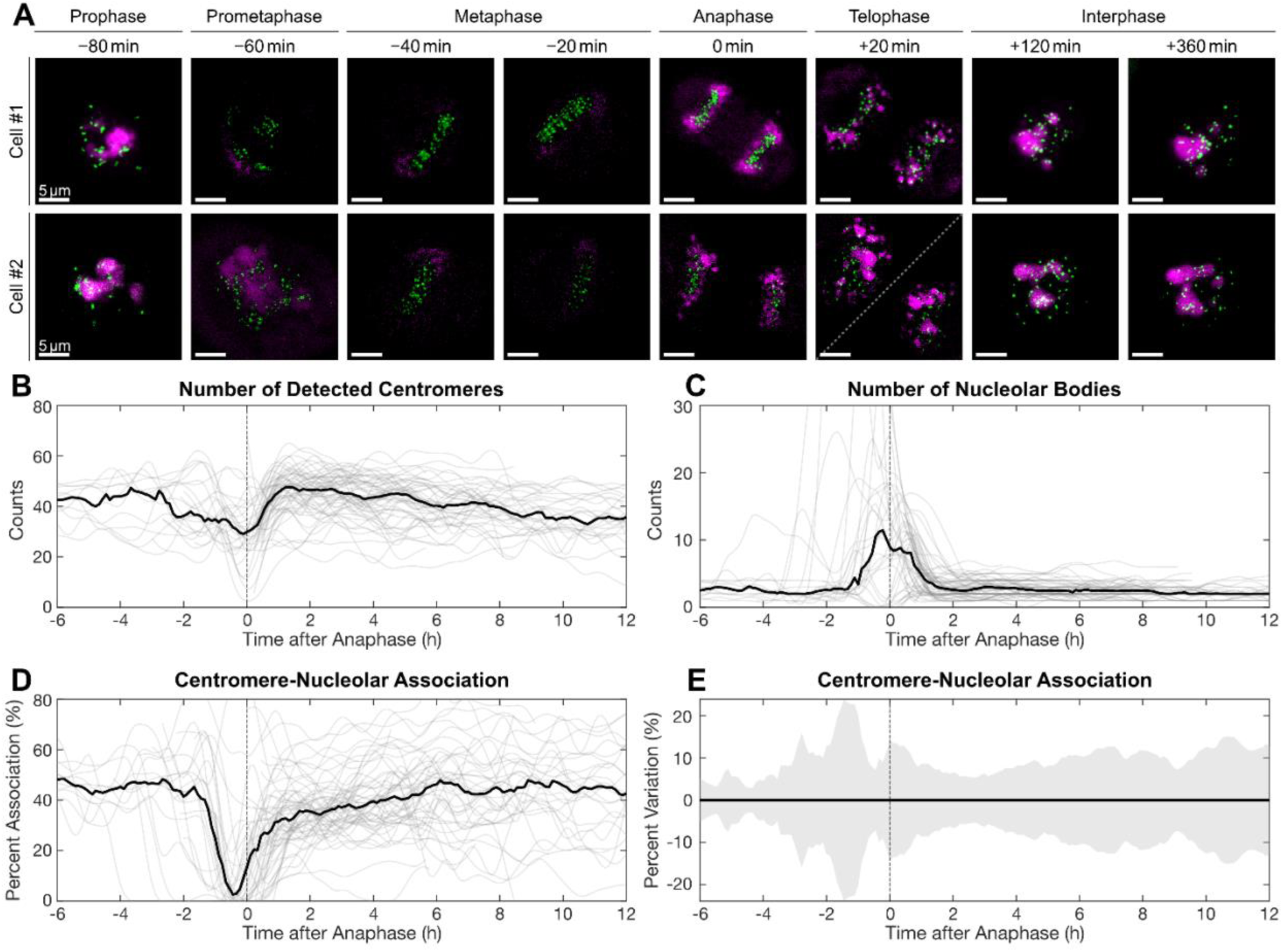
Centromere-nucleolus association during mitosis. **A.** Panel of images illustrating the dissociation of centromeres from nucleoli during mitosis and reassociation post-mitotically. **B.** Plots of the number of detected centromeres, **C.** number of nucleolar bodies, and **D.** centromere-nucleolar association from 6 hours before anaphase to 12 hours after anaphase. **E.** Illustration of the percent variation of centromere-nucleolar association from the median, from 6 hours before anaphase to 12 hours after anaphase.

**Figure 5.**
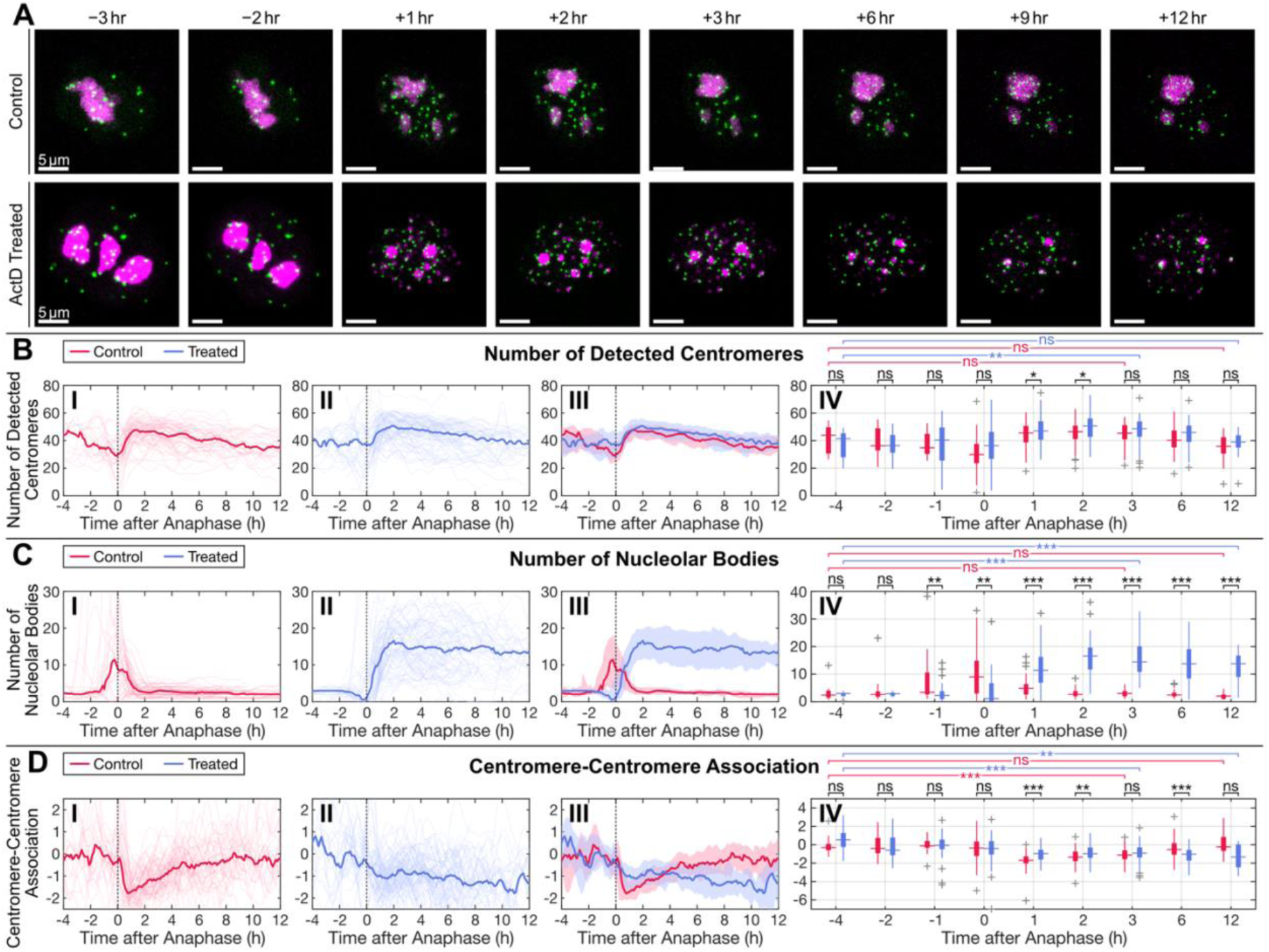
Effects of ActD treatment on centromere and nucleolar dynamics during mitosis. **A.** Panel of images illustrating the movement of centromeres relative to NPM1 from 3 hours before mitosis to 12 hours after mitosis. **B.** Plots of number of detected centromeres, **C.** number of nucleolar bodies, and **D.** centromere-centromere association, each illustrating **I.** smoothed raw data for control cells, **II.** smoothed raw data for ActD treated cells, **III.** an overlay of the control and treated cells, and **IV.** boxplots at different time points.

**Figure 6.**
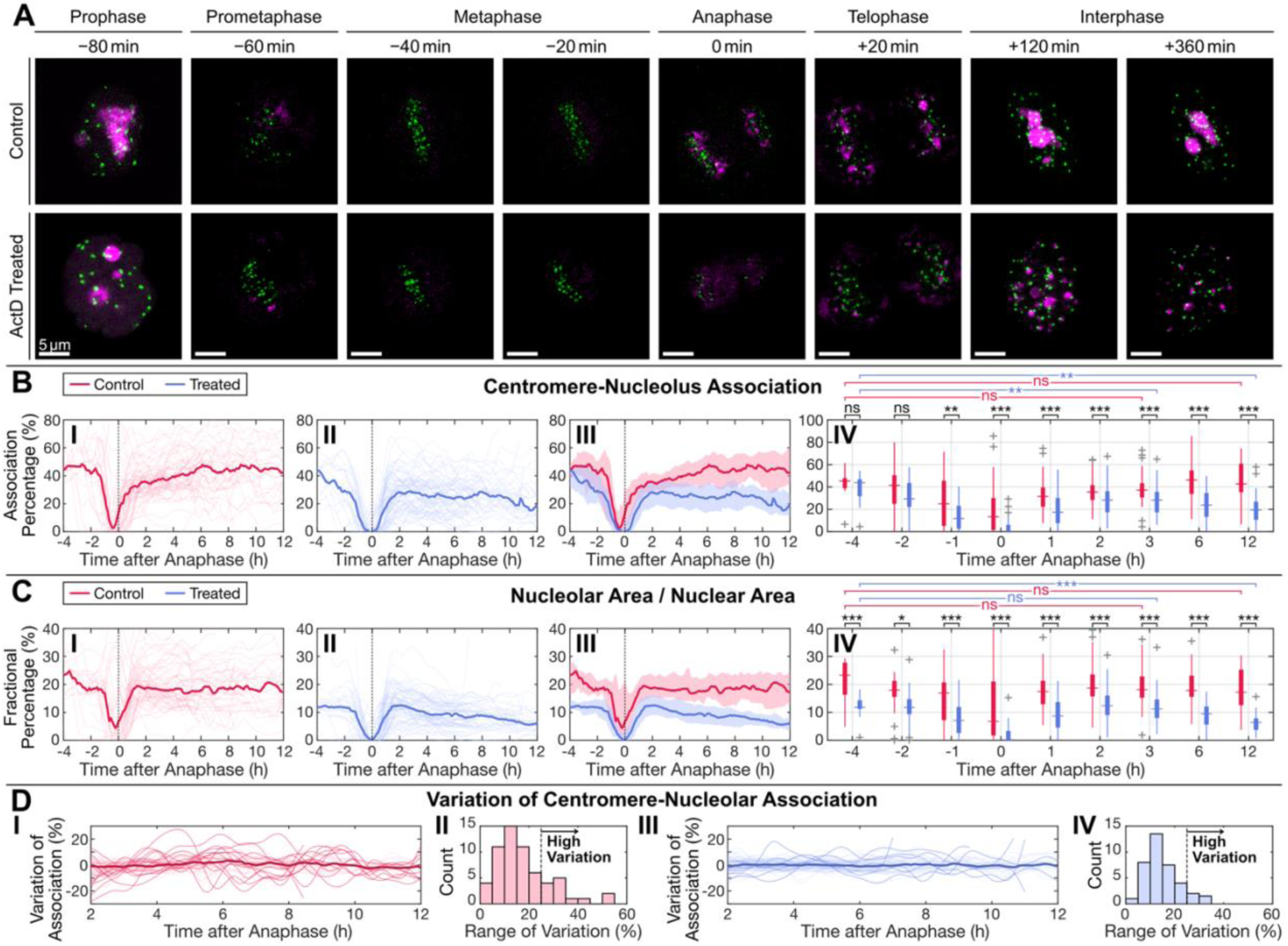
ActD treatment disrupts centromere-nucleolus associations. **A.** Panel of images illustrating the dissociation of centromeres from nucleoli during mitosis and reassociation post-mitotically. **B.** Plots of centromere-nucleolus association, and **C.** nucleolar area / nuclear area, each illustrating **I.** smoothed raw data for control cells, **II.** smoothed raw data for ActD treated cells, **III.** an overlay of the control and treated, and **IV.** boxplots at different time points. **D.** Plots of variation of centromere-nucleolar association, illustrating **I.** variation of association for control cells, **II.** histogram of range of variation for control cells, **III.** variation of association for treated cells, and **IV.** histogram of range of variation for treated cells.

To quantify these interactions, we first measured the number of nucleoli per cell over time (Fig. 3B) and applied the same trend-residual decomposition method as previously described. The results indicated no drastic changes in nucleolar body counts, with residuals fluctuating by approximately ±1, suggesting minor variations rather than systematic trends. We analyzed centromere-nucleolus associations by measuring the percentage of centromeres in close proximity to (touching), or overlapping with, nucleolar bodies in maximally projected image z stacks and applying the same decomposition method (Fig. 3C). The 2D measurement of the 3D centromere-nucleoli interaction is representative of the true interactions since nucleoli are the bulkiest structure in z and centromeres detected above or below nucleoli are unlikely not touching nucleoli. In HeLa cells (*n* = 60), centromere-nucleolus associations fluctuated over time, confirming that centromere positioning relative to nucleoli is dynamic during interphase. A subset of cells (13 out of 60) exhibited more pronounced fluctuations, with centromere-nucleolus association levels varying by more than 25% over time (Fig. 3C), indicating substantial spatial reorganization in these cells. We next examined whether variability in the number of detected centromeres per cell might influence the extent of centromere-nucleolus association (Fig. 3D). Using Spearman rank correlation, we found a value of −0.1011, indicating no meaningful correlation. Thus, the observed increase in nucleolar-centromere associations over time (Fig. 3C) is unlikely to be driven by fluctuations in centromere detection count, which may slightly decrease during extended imaging.

To further characterize the heterogeneity in centromere-nucleolus interactions per cell, we stratified the data from Fig. 3CIII into two groups of cells based on the extent of variation. The result is shown in Fig. 3E: low- (Fig. 3EI) and high-variation cells (Fig. 3EII). The range of variation was defined as the difference between the maximum and minimum values in each cell sample. This revealed a distribution of centromere-nucleolus association dynamics across the cell population, highlighting a subset of cells with significantly greater fluctuations (Fig. 3EIII). Since the observed variation cannot be due to the variation in the number of detected centromeres (Fig. 3D), it likely reflects the inherent dynamic nature of nucleoli-centromere interactions, capturing the real-time fluctuations in their spatial associations.

These findings demonstrate that while centromeres are commonly found at nucleolar periphery, some of their associations are dynamic, with individual centromeres exhibiting varying degrees of interaction with nucleoli over time. The variability in centromere clustering suggests potential functional implications in nuclear organization perhaps at different stages of cell cycle.

### Reestablishment of centromere clustering around nucleoli as cells enter G1

Live-cell imaging captured the stepwise dissociation and reorganization of centromeres relative to nucleoli throughout mitosis (Fig. 4A). At prophase (80 min), nucleoli remained intact. By prometaphase (60 min), nucleolar structures were largely disassembled, with only weak signals detectable, while centromeres grouped as chromosomes condensed into metaphase. During metaphase (40 to −20 min), nucleolar NPM1-mCherry signals were minimal, with faint remnants visible at the periphery of the metaphase plate. As cells progressed into anaphase (0 min), NPM1-mCherry began clustering at the edges of the separating chromosome bundles, marking the transition into two distinct daughter nuclei. At telophase (20 min), the association of pre-nucleolar bodies (NPM1-mCherry) with centromeres began (Fig. 4A), which most likely reflects the transcription activation at NORs, each forms a pre-nucleolar body. The percentage of centromeres associated with NPM1 bodies was not fully restored (Fig. 4D). As G1 progressed, the fusion of the pre-nucleolar bodies formed typical nucleoli (120 and 360 min) and nucleoli-centromere association closely resembled the pre-mitotic state.

To quantify the dynamic changes in nucleolar and centromere organization through mitosis, we analyzed several key parameters over time. The number of observed centromeres exhibited temporal fluctuations, remaining relatively stable before mitosis, ranging from ∼45 at −4 hours to ∼40 at −2 hours, before decreasing to ∼30 at 0 hours, likely due to mitotic condensation and overlapping centromeres (Fig. 4B). The number of detectable centromeres increased to ∼50 by 1 to 2 hours post-mitosis, before gradually declining again, likely due to a combination of progressive centromere clustering and photobleaching, which reduces the ability to resolve individual centromeres.

The nucleolar number as represented by the NPM1 labeling, gradually increased from an average of 2 to 3 per nucleus at −4 hours to ∼5 at −1 hour as chromosome condensation proceeded and nucleoli broke down. The number sharply increased to ∼10 clustered labeling at 0 hours, corresponding to anaphase (Fig. 4A and 4C). The higher number of NPM1 bodies at late mitotic phase and early G1 phase indicate the association of NPM1 with mitotic chromosomes and nucleating towards active NORs to form pre-nucleolar bodies on each of the acrocentric chromosomes. This number then gradually decreased to 4 to 5 nucleoli at 1 hour and returned to baseline levels (2 to 3 per nucleus) by 2 hours post-anaphase, reflecting the fusion of pre-nucleolar bodies to form typical nucleoli as new daughter cells formed. Interestingly, NPM1 becomes associated with mitotic chromosomes at anaphase with enrichment at both ends of the parting chromosome line-ups (Fig. 4A). The functional significance of the interactions is unclear.

To quantify the dynamics of centromere-nucleolus interactions through mitosis, we measured the fraction of centromere associated with NPM1 signals over time. Prior to mitosis, the centromere-nucleolus association was relatively stable at 45% to 50% from −4 to −2 hours, but it dropped to 0% at metaphase (−40 to −20 minutes) (Fig. 4A and D) and began to increase to ∼5% at 0 hours (anaphase), coinciding with nucleolar disassembly (Fig. 4D). As nucleoli began to reassemble, the fraction of associated centromeres gradually increased to ∼20% at 0 hour, ∼30% at 1 hour, and ∼35% at 2 hours, before fully recovering to pre-mitotic levels (40% to 50%) approximately 6 hours post-mitosis. These trends suggest that centromere-nucleolus interactions are actively disrupted during mitosis as nucleoli disassemble and gradually restored as cells transition into G1 phase as nucleoli reform.

These findings demonstrate that centromere-nucleolus associations undergo a dynamic dissociation-reassociation cycle across mitosis and early G1. Centromeres disengage from nucleoli during mitotic progression and re-establish clustering post-mitotically in a time-dependent manner, highlighting the interplay between centromeres and nucleoli in nuclear reorganization.

### ActD treatment alters interphase nucleoli and centromere organization and dynamics

As previously reported, rDNA transcription is important for nucleoli integrity. We used a low concentration of ActD treatment at 0.04 µg/mL to evaluate whether the changes of nucleolar structure due to the loss of rDNA transcription could affect centromere and nucleolar dynamics during interphase. We compared live-cell imaging data of vehicle-treated and ActD-treated cells. The results revealed striking differences in nucleolar organization and centromere dynamics upon ActD treatment (Fig. 5A). In control cells, nucleolar bodies exhibited clustering and dynamic reorganization, whereas in ActD-treated cells, nucleoli became condensed prior to mitosis and were unable to coalesce to form typical nucleoli after mitosis. Additionally, centromeres in treated cells appeared much more stationary, suggesting a restriction in their mobility compared to controls.

ActD treatment did not affect centromere numbers between treated and control untreated cells (Fig. 5B), which likely decreased over time due to progressive centromere clustering or imaging-related factors such as photobleaching. In contrast, the number of nucleoli was drastically altered in ActD-treated cells, but only after mitosis (Fig. 5C) cells. Control cells maintained ∼2 ± 0.5 nucleolar bodies per nucleus (Fig. 5CI), apart from transient increases caused by mitotic clustering around anaphase chromosomes. ActD-treated cells showed no significant change in nucleoli number prior to mitosis (Fig. 5CII); however, following mitosis, they exhibited a striking increase, with ∼17 ± 4 nucleolar bodies per cell after mitosis (Fig. 5CII and III). Statistical comparisons at all post-mitotic time points indicated highly significant differences (*** in Fig. 5CIV), confirming that ActD profoundly alters nucleolar organization in post-mitotic cells. Analysis of the total nucleolar area revealed no decrease in overall NPM1 levels (Supp. Fig. S7A), indicating that despite NPM1 remaining abundant, centromeres do not associate with it. Consistent with this, the characteristic clustering of NPM1 around anaphase chromosomes observed in controls was also absent in ActD-treated cells (Fig. 5CII and III), indicating NPM1 associations with chromosomes were disrupted.

ActD treatment also altered centromere-centromere associations compared to untreated cells (Fig. 5D). In control cells (Fig. 5DI), centromere-centromere association values were ∼0 prior to mitosis (indicating random distribution), transiently decreased to −1.6 after mitosis (indicating centromere dispersion), and returned to baseline (∼0) within ∼8 hours. By contrast, ActD-treated cells (Fig. 5DII) exhibited higher association values (∼1) prior to mitosis, followed by a progressive decrease to −1.9 over ∼9 hours post-mitosis, indicating persistent centromere dispersal. Direct comparison (Fig. 5DIII) revealed that while control cells re-established pre-mitotic centromere-centromere associations 12 hours post-mitosis compared to −4 hours prior, ActD-treated cells failed to recover. Consistent with this, statistical analysis showed no difference between pre- and post-mitotic associations in controls, whereas ActD-treated cells exhibited significant changes across the same interval. Importantly, quantification of total nucleolar area, overall nucleolar intensity, and nuclear area (Supp. Fig. S7A-C) confirmed that the overall NPM1 signal within the nucleus remained stable between ActD-treated and control cells, indicating that ActD alters nucleolar organization without reducing NPM1 levels.

These findings demonstrate that ActD treatment disrupts nucleolar organization, leading to decreases in variability in centromere-nucleolus interactions, thus the dynamics of centromeres to and from nucleoli. While total centromere counts remain unaffected, the disruption of nucleoli in treated cells likely contributes to the reduced stability of centromere-nucleolus associations. Additionally, the reduced centromere mobility observed in ActD-treated cells suggests that nucleolar integrity or the effect of DNA intercalation of ActD may play roles in regulating centromere positioning and movement in interphase cells.

### ActD treatment disrupts nucleolar reassembly post-mitotically and nucleoli-centromere interactions

Live-cell imaging of control and ActD-treated cells captured the progression of nucleoli and centromeres through mitosis (Fig. 6A). In control cells, nucleolar numbers averaged ∼2.4 per nucleus (Fig. 3B) and NPM1 displayed the characteristic concentrated labeling around mitotic chromsomes during anaphase. In ActD-treated cells, nucleolar numbers are similar to those of the control prior to mitosis. Strikingly, upon anaphase, ActD-treated cells failed to show the typical clustering of NPM1 around chromosomes. As ActD treated cells entered G1, pre-nucleolar bodies were unable to fuse into mature nucleoli, persisting instead as numerous smaller foci (Fig. 5A and 6A). This defect resulted in a substantial post-mitotic increase in nucleolar numbers (Fig. 5CII, III, IV), highlighting that inhibition of Pol I transcription prevents normal nucleolar reassembly and disrupts nucleolar structure.

To assess how ActD influences centromere-nucleolus interactions, we measured the fraction of centromere associated with nucleoli over time (Fig. 6A and B). We show the centromere-nucleoli association is reduced in ActD-treated cells compared with control cells (Fig. 6B). In addition, we observed a different behavior in ActD-treated cells when we compare the centromere-nucleolus interactions between pre- and post-mitotic cells. While control cells showed no significant changes in centromere-nucleolus association pre- vs post-mitosis, ActD-treated cells exhibited a marked reduction in centromere-nucleolus association. In addition, the nucleolar areas are also significantly reduced upon treatment with ActD before and after mitosis (Fig. 6C), consistent with previous findings of nucleoli segregation and condensation when treated with ActD (Snow, 1972; Shav-Tal et al., 2005).

To evaluate the effect of ActD treatment on centromere-nucleolus interactions, we analyzed the variation in nucleolar-centromere association over time (Fig. 6D) using the same trend-residual decomposition method as in Fig. 2C, extracting the residuals to quantify fluctuations. Figure 6DI displays residuals for control cells, while Fig. 6DIII the bottom panel presents residuals for ActD-treated cells, with accompanying histograms illustrating the distribution of variation ranges in each condition. In control cells, 13 out of 60 cells exhibited high variation in centromere-nucleolus interactions (Fig. 6DII), whereas in ActD-treated cells, only 7 out of 75 cells displayed similarly high variation (Fig. 6DIV), defined as a range of centromere-nucleolus association fluctuations exceeding 0.25. Statistical analysis using a Fisher’s exact test yielded a *p*-value of 0.0394, indicating a significant reduction in the proportion of cells with more dynamic nucleoli-centromere interactions upon ActD treatment. These results suggest that ActD treatment stabilizes centromere-nucleolus interactions, reducing their variability and potentially restricting centromere repositioning relative to nucleoli.

These findings demonstrate that ActD treatment disrupts nucleolar integrity, leading to nucleolar segregation, reduced nucleolar area, reduced centromere-nucleolus interactions, and decreased NPM1 association with mitotic chromosomes. As ActD selectively inhibits rDNA transcription at this concentration, these data are consistent with published studies where the loss of Pol I activity disrupts nucleolar structure and nucleoli-centromere interactions.

## DISCUSSION

We analyzed the intra-nuclear organization of centromeres and their relationship with nucleoli. With GFP-CENPA marking centromeres and mCherry-NPM1 labeling nucleoli, we recorded the dynamics of centromeres and nucleoli-centromere interactions in living cells over 10 hours with or without the treatment of ActD that selectively inhibits rDNA transcription and disrupts nucleolar structure. The results show some unexpected fast movement of a subset of centromeres in nuclei and a fluid relationship between centromeres and nucleoli, which are substantially altered when cells were treated with a low concentration of ActD.

### Centromere dynamics

In this study, we visualize and quantitatively analyze interphase centromere behavior in living HeLa cells by expressing GFP–CENP-A. Although GFP–CENP-A does not fully recapitulate the biochemical properties of endogenous CENP-A at the nucleosome level (Kalitsis et al., 2003), the tagged protein coexists with endogenous CENP-A in our HeLa cells, and the stable expression does not measurably perturb cell proliferation or mitotic process. Thus, GFP–CENP-A is suitable for monitoring centromere dynamics without detectable adverse effects on HeLa cell viability.

Time-lapse imaging revealed that a subset of centromeres exhibits substantial mobility within interphase nuclei, with displacements of up to several micrometers over the course of hours (Fig. 2A). These movements cannot be attributed to nuclear rotation, as centromeres moved in divergent directions within the same nucleus and nucleolar positioning remained largely constant (Fig. 2A; Suppl. Fig. 2). We also observed centromere fusion events, in which two or more foci transiently come together (Fig. 1D–E; Fig. S1). Whether these represent interactions between homologous centromeres or heterologous associations is unclear.

Given that centromeres are embedded within extensive pericentromeric heterochromatin—typically positioned at the interior of chromosome territories—it is intriguing to consider the functional implications of their mobility. We classify centromere movement into 2 distinct classes. First, centromere movement may reflect directed interactions with nucleoli, consistent with observed trajectories toward or away from nucleolar structures (Fig. 3A; Suppl. Fig. 6). Second, centromere motion lacking an obvious directional target may indicate dynamic changes in heterochromatin organization or broader remodeling of the specific chromosome territories.

### Nucleolar associated centromere dynamics

Centromere (particularly pericentromeric heterochromatin) nucleolar interactions represent a prominent feature of interphase nuclear organization (Németh et al., 2010; van Koningsbruggen et al., 2010; Bersaglieri and Santoro, 2019; Bersaglieri et al., 2022). Although the rDNA arrays on acrocentric chromosomes are expected to associate with nucleoli, a substantially larger fraction of centromeres localizes to nucleoli, indicating a broader organization mechanism. These spatial interactions vary through development and differentiation stages: they are highly enriched in pluripotent stem cells, diminished with differentiation, and elevated again in transformed cancer cells (Rodrigues et al., 2022), suggesting that nucleolar tethering of centromere reflects cell-state–dependent chromatin architecture and spatial organization.

This study in living HeLa cells shows that the nucleoli-centromere interactions are intrinsically dynamic. Centromeres rapidly re-establish nucleolar contact as cells enter early G1, coincident with nucleolar reassembly, and the extent of association continues to increase through G1. Individual cells exhibit marked fluctuations in the proportion of nucleolus-associated centromeres (Fig. 3E), reflecting true positional changes rather than differences in signal detection. These dynamics correlate with micrometer-scale centromere mobility, transient fusion events, and oscillatory movements toward and away from nucleoli.

Could centromere associations with nucleoli be related to the assembly of centromeres? Some of the key factors involved in centromere assembly are nucleolar proteins. NPM1 is a nucleolar protein and a known components of the CENP-A pre nucleosome complex (Dunleavy et al., 2009; Foltz et al., 2009). In addition, the CENP-A specific histone chaperone HJURP that is necessary for CENP-A deposition to centromeres in G1 is transiently associated with the nucleolus in the S-phase. However, not all centromeres are associated with nucleoli in G1-phase and a clear mechanistic link between centromere-nucleolar association and centromere assembly has not yet been established.

Nearly all pericentric heterochromatins have been shown to be associated with nucleoli by DNA sequencing of isolated nucleoli (van Koningsbruggen et al., 2010; Bersaglieri and Santoro, 2019; Vertii et al., 2019; Bizhanova et al., 2020; Bizhanova and Kaufman, 2021; Bersaglieri et al., 2022; Peng et al., 2023). Interestingly, a recent study showed that not all of them are associated with nucleoli equivalently; rather, pericentric heterochromatins can be classified into distinct groups (G1, G2, and G3) based on their interaction patterns and proximity to centromeres, with G2 hNADs (high-confidence nucleolus-associated domains) located closest to centromeres of NOR containing chromosomes with strongest cis-chromosome-chromosome interactions, while G3 hNADs show stronger inter-chromosal interactions with higher levels trans interactions (Peng et al., 2023). The sptatial and temporal differential associations to nucleoli by different pericentric heterochromatins are consistent with the observed movement of centromeres from and towards nucleoli.

The question is then whether centromeres associate with nucleoli through intrinsic properties of the centromere itself, or whether they are carried to nucleoli as passengers via pericentromeric heterochromatin that directly interacts with nucleolar components. Our unpublished observations support the latter model. Specifically, an artificial neocentromere containing a CENP-A–rich domain but lacking flanking pericentromeric heterochromatin fails to associate with nucleoli (unpublished data). On the other hand, the findings that human ES cells have the maximal centromere clustering around nucleoli (over 70%) (Rodrigues et al., 2023) with minimal heterochromatin state points towards the less significant role of pericentromeric heterochromatin in anchoring centromeres to nucleoli. This point opens up another possibility that the anchoring centromeres to nucleoli can be a mechanism for optimal chromosome spatial organization that controls the genome expression through spatial physical constraints catering to the specific states of cells. The successful testing of this hypothesis could help shed light into the logic and practical functionality of spatial genome organization and how this organization governs genome expression.

### Centromere movement unrelated to nucleoli

What drives the movement of centromeres in nucleoplasm, sometimes over µm distance (Figs. 2AIII, 2AVI)? One possible explanation could be due to the activation or inactivation of genes near centromeres which drag the loci towards specific direction for optimal expression. Future studies will attempt to address the dynamics of specific pericentromeric loci, particularly for centromeres associated with genes whose expressions change through cell cycle. For example, some gene activation could lead to direct movement of the loci towards nuclear speckles in micrometers (Khanna et al., 2014).

Conversely, another possibility exists that centromeres could help reposition specific chromosomes in response to cellular activity cues. They could represent changes of the heterochromatin state or the reorganization of the chromosome territories. Recent studies indicate that distinct pericentromeric heterochromatin domains associate with nucleoli at specific cell-cycle stages (Peng et al., 2023), providing a compelling mechanistic basis for the directional movements we observe. Transcriptional activity near centromeres may further bias locus positioning. Together, our findings support a model in which centromere mobility integrates cell-cycle cues, heterochromatin state, and nucleolar dynamics to modulate higher-order genome organization.

### Alterations of Centromere-nucleoli interactions in the presence of ActD

Treatment with a low concentration of ActD selectively inhibits Pol I transcription by preferentially alkylating GC rich DNA. Many studies showed that ActD treatment induces nucleolar segregation where the rDNA and fibrillar components reorganize towards the edge of the nucleoli forming cap like structures (Shav-Tal et al., 2005). Furthermore, the reduction of Pol I polymerase through siRNA or degron also showed similar distortion of nucleoli (Wang et al., 2023). Along with the changes of nucleolar structure, the centromere association reduces (Wang et al., 2023). Our observations in the live cell imaging are consistent with these findings that the clustering of centromeres around nucleoli reduced in the presence of ActD (Fig. 6B). In addition, dynamics of centromeres also significantly reduced (Fig. 6D). Could it be the Pol I transcription inhibition or DNA structural integrity that impact the dynamic of centromere? In our previously published study of chromosome dynamics in living cells with the smallest area of photoactivation of H3-Dendra, treatment with ActD at the same concentration substantially reduced overall chromosome movement. Thus, the alkylation of GC rich DNA could be a factor in centromere dynamic reduction, which could prevent them from approaching nucleoli before mitosis and pre-nucleolar bodies post-mitotically as well as intranuclear movement. In addition, the disruption of nucleolar structure could also play a role in preventing centromere attachment. Perhaps some of the centromeres are no longer affiliated with nucleoli due to its surface changes.

Interestingly, the live cell studies using NPM1 labeling, nucleoli show some unique features of the multi-functional nucleolar protein. NPM1 as part of the granular components does not show segregated morphology when treated with ActD as observed for fibrillar proteins, including rDNA transcription factors and some of the pre-rRNA processing factors which move to the periphery of the distorted nucleoli forming cap or dot-like structures. It only becomes rounder and condensed pattern after exposure to ActD. In untreated cells, NPM1 dissociates as cells enter mitosis along with the disassembly of nucleoli, but becomes clustered around both ends of segregated chromosome at early anaphase and expanding around the still condense chromosomes as cytokinesis commences (Fig. 4A).

In summary, with red nucleoli and green centromeres in living HeLa cells, we observed dynamic movement and fusion of centromeres through different phases of the cell cycle. Some centromeres in interphase HeLa cells show motions towards and away from nucleoli, while other not associated with nucleoli move directly through nucleoplasm. The nucleolar association of centromeres generally breaks down as cells enter mitosis and re-establishes as cells progress into G1. Treatment with ActD, substantially restricted the movement and nucleolar association. Centromeres and their association with nucleoli are dynamic and oscillate through different phases of the cell cycle. Does the centromere dynamic represent a way of spatial genome organization regulation and its impact on genome expression remain questions for future investigations.

## ACKNOWLEDGEMENTS

We thank the support from the National Institutes of Health (R01GM139151, R01GM140478, and U54CA268084 to HFZ, U01CA260699 to DF and SH), the Christina Enroth-Cugell Graduate Research Award at Northwestern University for WY. We also thank Drs. João Mamede and Tom Hope, as well as the Center for Advanced Microscopy at Northwestern University’s Feinberg School of Medicine, for their microscopy assistance.

## Data Availability Statement

Data available on request from the authors.

