## Supplemental Document for "The dynamics of nucleolus-centromeres interaction in living cells"

### Supplemental Note:
### The dynamics of nucleolus-centromeres interaction in living cells


Wei-Hong Yeo[a], Emma Freeman[b], Alex B. Willis[c], Hao F. Zhang[a], Daniel Foltz[c], and Sui Huang[b,*]

[a] Department of Biomedical Engineering, Northwestern University, Evanston, IL, USA
[b] Department of Cell and Developmental Biology, Northwestern University, Chicago, IL, USA
[c] Department of Biochemistry and Molecular Genetics, Northwestern University, Chicago, IL, USA

*Corresponding Author: Sui Huang

Department of Cell and Developmental Biology

Northwestern University

Feinberg School of Medicine

303 E. Chicago Ave. Searle building, Rm 6-450

Chicago, IL, 60614, USA

312-503-4269

#### SUPPLEMENTARY NOTE 1: RIPLEY'S *K* AND *L* FUNCTIONS

##### *Uncorrected Ripley's K and L Functions*

Ripley's *K*-function provides a non-parametric summary of point-to-point interactions in two dimensions. Given a point pattern of $N$ points in a region of true area $A$, with density $\lambda = N/A$, the uncorrected estimator at scale $r$ is:

$$\widehat{K}(r) = \frac{\lambda^{-1}}{N} \sum_{i \neq j} \mathbb{1}\big(d_{ij} \leq r\big), \tag{1}$$

where $d_{ij}$ is the Euclidean distance between points $i$ and $j$, $\mathbb{1}(\cdot)$ is the indicator function, which gives 1 when the condition is true and 0 otherwise.

Under complete spatial randomness (CSR), the points follow a homogeneous Poisson process, and one has $\widehat{K}_{\text{CSR}}(r) = \pi r^2$, so that departures of $\widehat{K}(r)$ above or below $\pi r^2$ indicate clustering or inhibition, respectively.

For data analysis, the variance-stabilized *L*-function is more commonly used instead:

$$\hat{L}(r) = \sqrt{\frac{\widehat{K}(r)}{\pi}}, \tag{2}$$

which, under CSR, satisfies $\hat{L}(r) = r$. Therefore, when one plots $\hat{L}(r) - r$, the deviations above zero signify clustering at scale $r$ (more neighbors than expected), whereas deviations below zero indicate inhibition or regularity (fewer neighbors than expected). Because the square-root transformation linearizes the quadratic growth of $\widehat{K}(r)$, the variance of $\hat{L}(r)$ is more nearly constant across $r$, making it easier to identify scales of interaction.

Supp. Fig. S1 illustrates these concepts with three synthetic point patterns – regular, random, and clustered – each overlaid by concentric circles at radii 5, 10, and 15 pixels and a sprinkling of uniform "noise" points. In the regular pattern, $\hat{L}(r) - r$ falls below zero for $r < 20$, reflecting the enforced spacing of points; in the Poisson pattern it hovers around zero; and in the clustered model it rises above zero, peaking at $r = 10$, illustrating that the points form a cluster with $r \approx 10$. This demonstrates the sensitivity of Ripley's *L*-function to different spatial organizations.

##### *Edge-Correction and Density Estimation*

In spatial point-pattern analysis with limited data, such as quantifying centromere organization, points counts are limited, and the observation window is irregular. This can bias both the estimated density and the pair-counts.

**Buffered Convex-Hull Area.** To mitigate boundary effects, we first define the observed set of points, $X = \{x_1, \ldots, x_n\}$ and $\text{convhull}(X)$ to be its convex hull. We can then denote its raw area by $A = |\text{convhull}(X)|$. This polygon provides a data-driven approximation of the sampling region, but if used directly in $\lambda = n/A$, it underestimates the true area accessible to the neighborhoods of radius $r$. In particular, the points at the hull boundary cannot "see" a full circle of radius $r$, leading a systematic undercounting of neighbors.

Therefore, to ensure that every disk of radius $r$ around an observed point lies entirely within the sampling window, we dilate the convex hull by $r$. We define the Minkowski sum

$$W_r = \{x : \exists y \in \mathrm{convhull}(X) \text{ with } \|x - y\| \leq r\}, \tag{3}$$

and let $\hat{A}(r) = |W_r|$. This serves as the effective region in which circles of radius [illegible] around any data point lie fully inside the window. The point-density estimate is then $\hat{\lambda}(r) = n/\hat{A}(r)$, which corrects for the fact that circles of radius $r$ centered at boundary points extend beyond the raw hull.

**Isotropic Edge-Correction Weights.** Points near the boundary have a truncated neighborhood, as a circle of radius $d_{ij}$ around point $i$ may protrude outside the window. To correct for this bias, each pair $(i, j)$ is assigned a weight

$$w_{ij} = \frac{1}{2\pi} \int_0^{2\pi} \mathbb{1}\left(x_i + d_{ij}[\cos\theta, \sin\theta] \in \mathrm{convhull}(X)\right) \, \mathrm{d}\theta, \tag{4}$$

i.e., the fraction of the full 360° circle around $x_i$ at radius $d_{ij}$ that lies inside the original convex hull.

##### ***Fully Corrected Estimator***

With these definitions, the unbiased estimator of the Ripley's *K*-function becomes

$$\hat{K}(r) = \frac{\left[\hat{\lambda}(r)\right]^{-1}}{N} \sum_{i \neq j} \frac{\mathbb{1}(d_{ij} < r)}{w_{ij}}. \tag{5}$$

Finally, combining Eqs. (2) and (5), this yields unbiased, variance-stabilized estimates of $\hat{L}(r) - r$ suitable for comparing spatial point patterns in irregular domains.

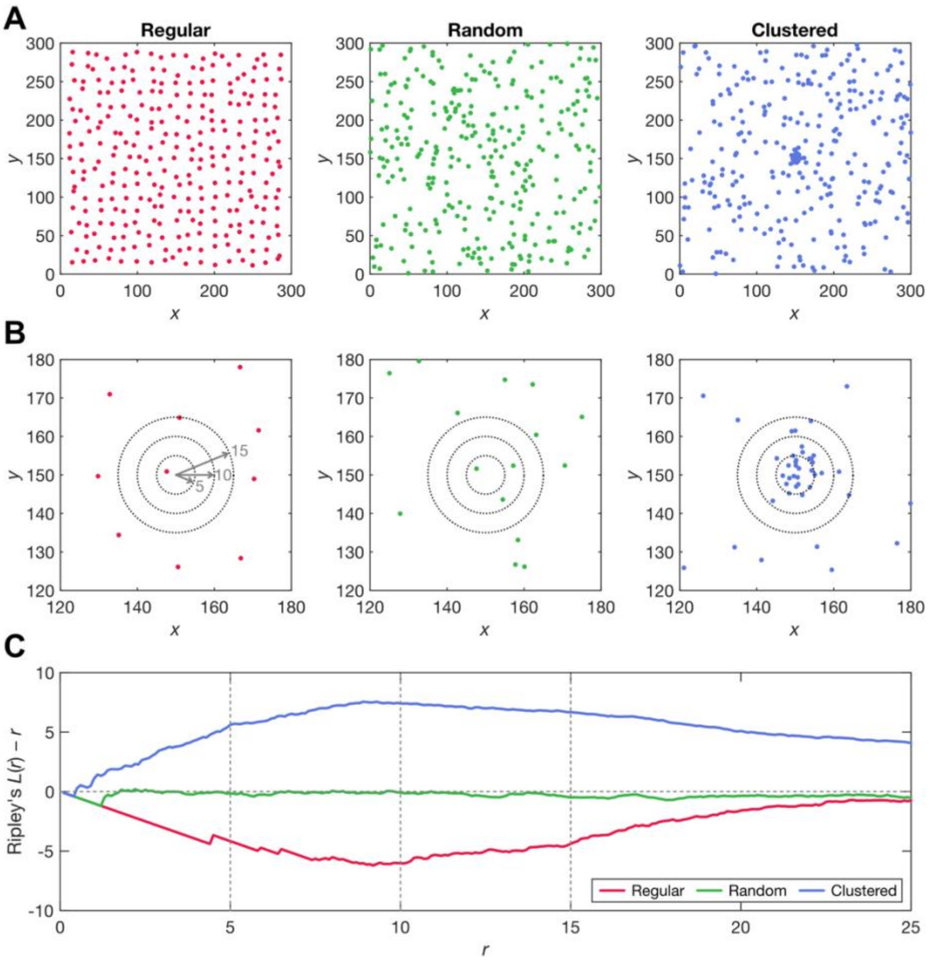


***Supplemental Figure 1. A.*** *Full-domain point distributions (300 x 300 units):* ***I.*** *jittered regular grid,* ***II.*** *homogeneous Poisson (random) pattern, and* ***III.*** *Neyman–Scott cluster process. Points are colored by pattern type.* ***B.*** *Zoomed-in region for each pattern, overlaid with concentric guide circles of radii 5, 10, and 15 units. These circles indicate the scales at which spatial clustering or regularity is assessed.* ***C.*** *Ripley's L(r) – r curves for each pattern, computed over r = 0.1–25 units. Vertical lines mark r = 5, 10, and 15 units. The regular pattern exhibits L(r) – r < 0 at small r (inhibition), the Poisson pattern remains near zero (randomness), and the clustered pattern shows L(r) – r > 0 (aggregation).*

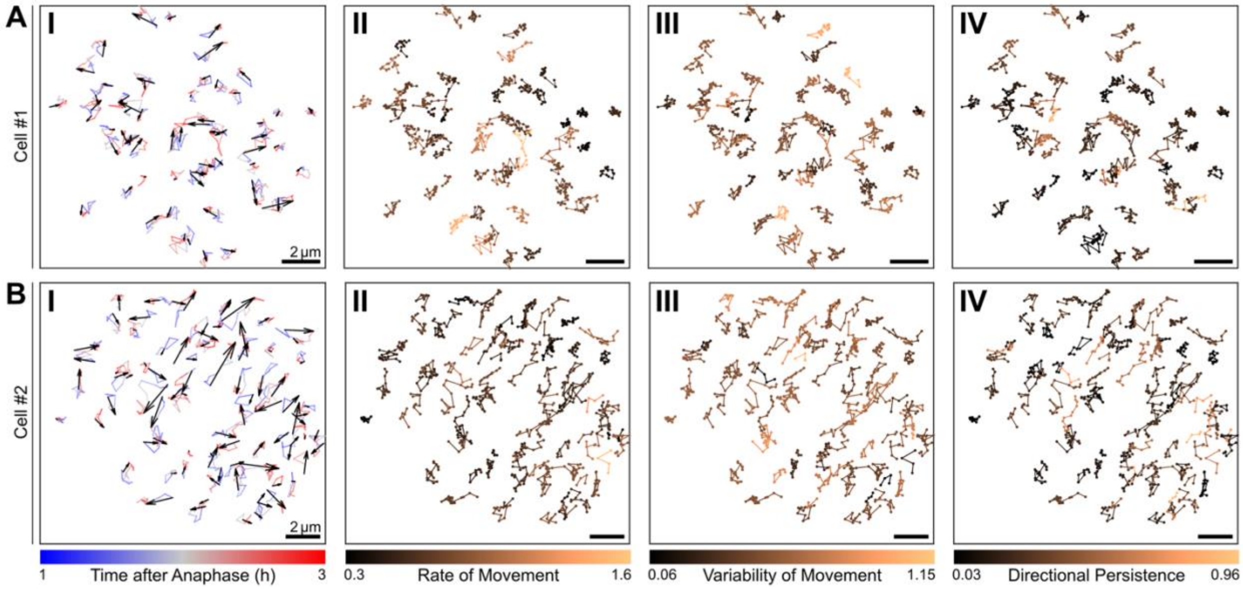


***Supplemental Figure 2*** *Plots of* ***I.*** *displacement of centromeres,* ***II.*** *rate of movement,* ***III.*** *variability of movement,* ***IV.*** *directional persistence for* ***A.*** *cell #1 and* ***B.*** *cell #2 of Fig. 2A.*

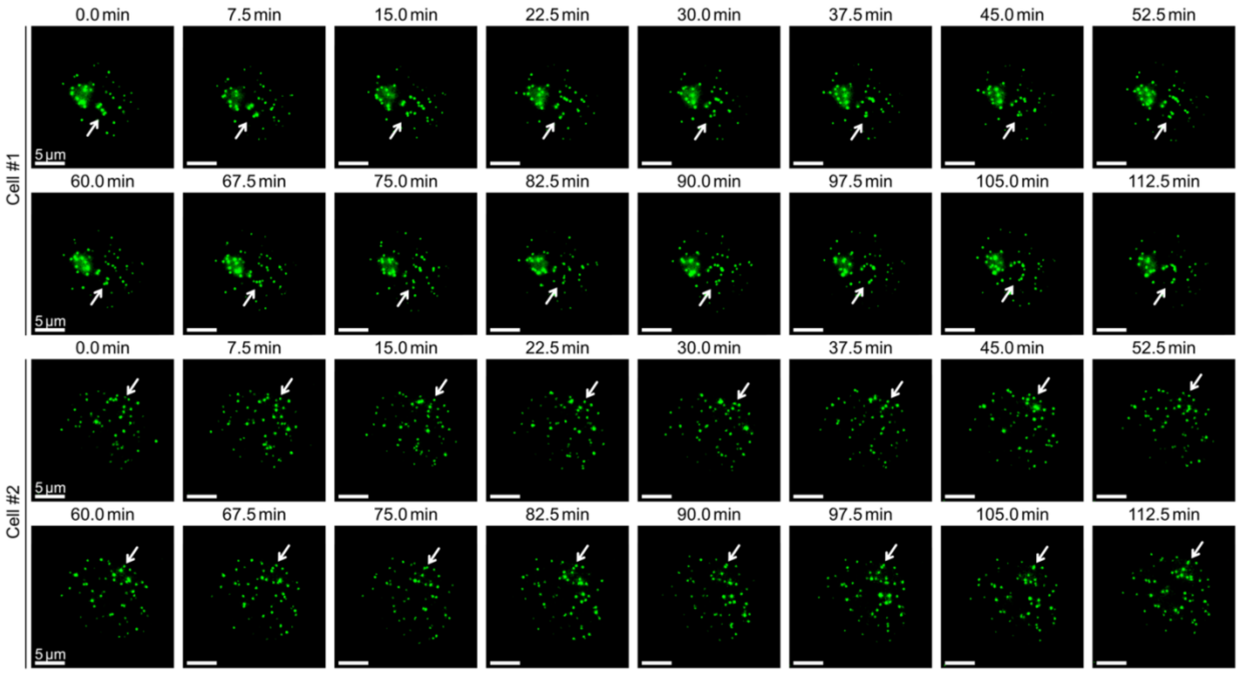


***Supplemental Figure 3*** *Panel of images illustrating the movement of centromeres over 2 hours.*

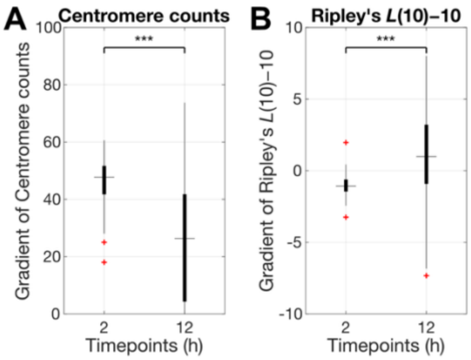


***Supplemental Figure 4. A.*** *Comparison of the gradient of centromere counts at 2 and 12 hours timepoints.* ***B.*** *Comparison of the gradient of Ripley's L(10)−10 at 2 and 12 hours timepoints.*

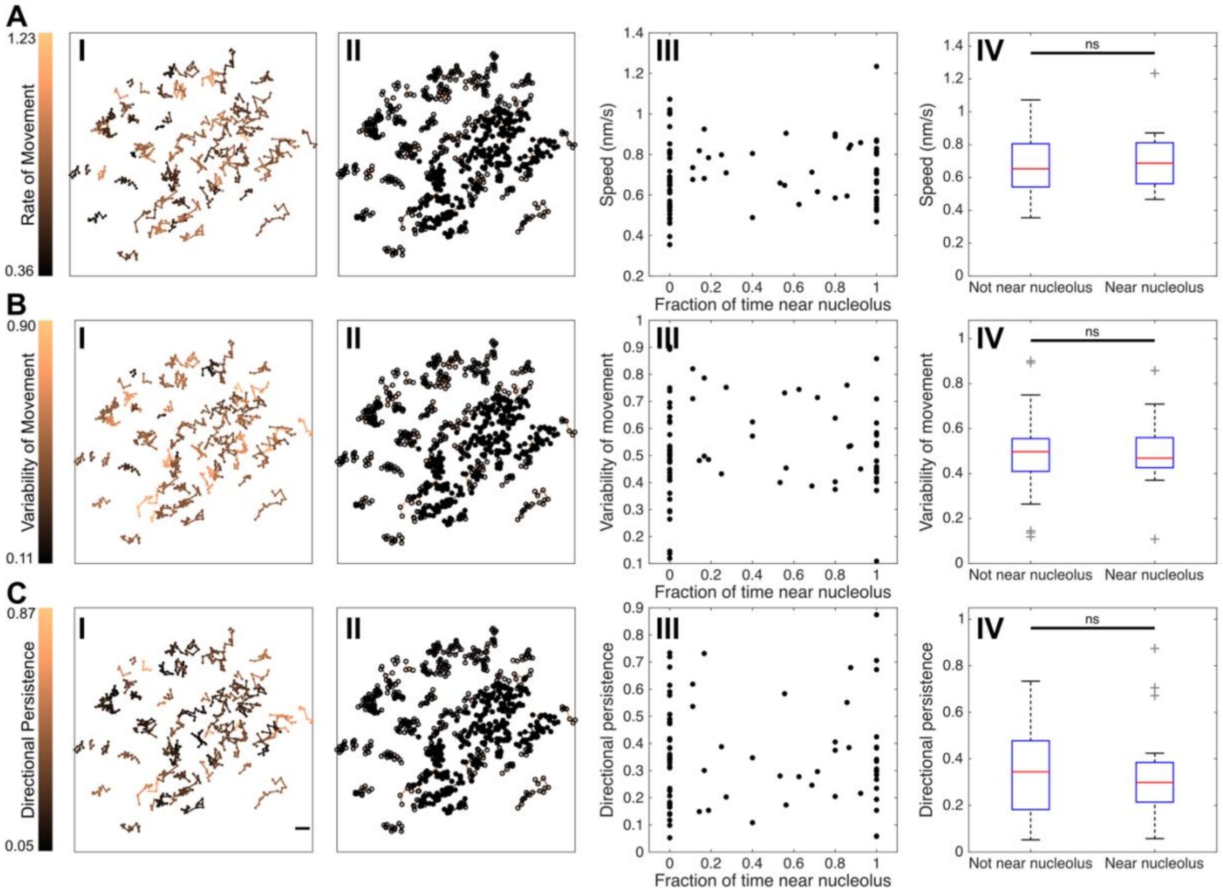


***Supplemental Figure 5*** *Plots of* ***A.*** *rate of movement,* ***B.*** *variability of movement, and* ***C.*** *directional persistence, each visualized as* ***I.*** *colored tracks in spatial coordinates,* ***II.*** *dots plotted when it is near a nucleolar body,* ***III.*** *plots against fraction of centromeres near a nucleolar body, and* ***IV.*** *boxplots comparing centromeres near and not near nucleolar bodies.*

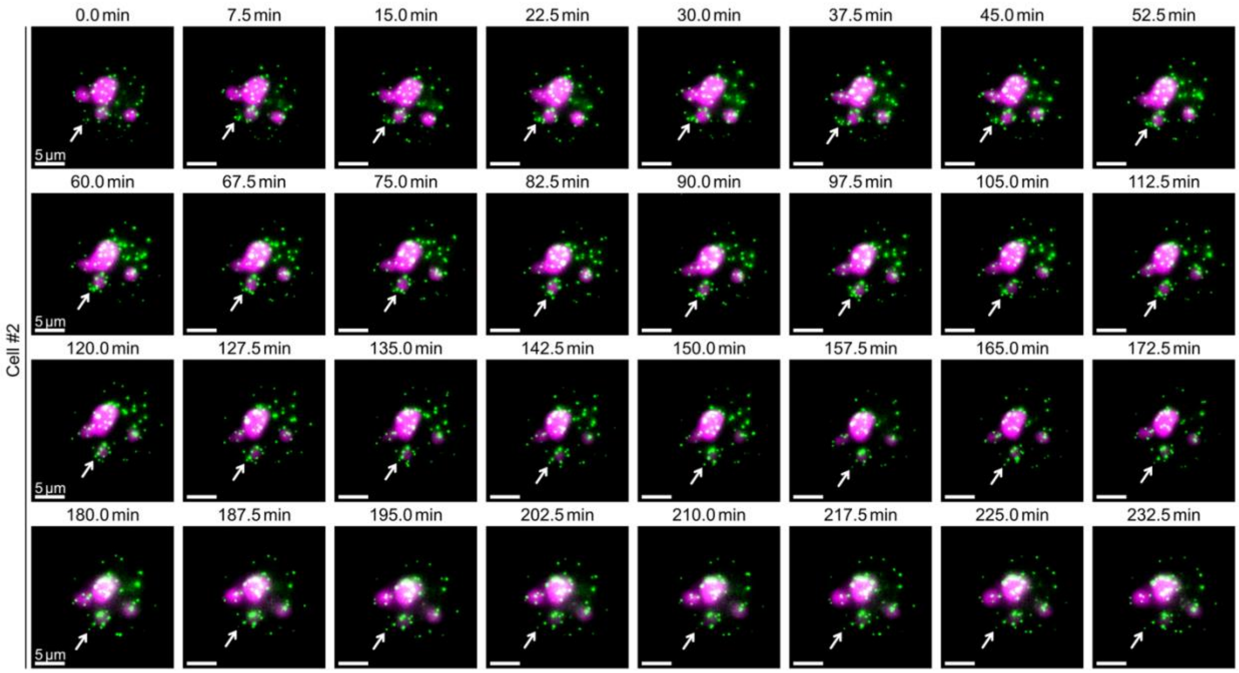


***Supplemental Figure 6*** *Panel of images illustrating the movement of centromeres relative to NPM1 over 4 hours.*

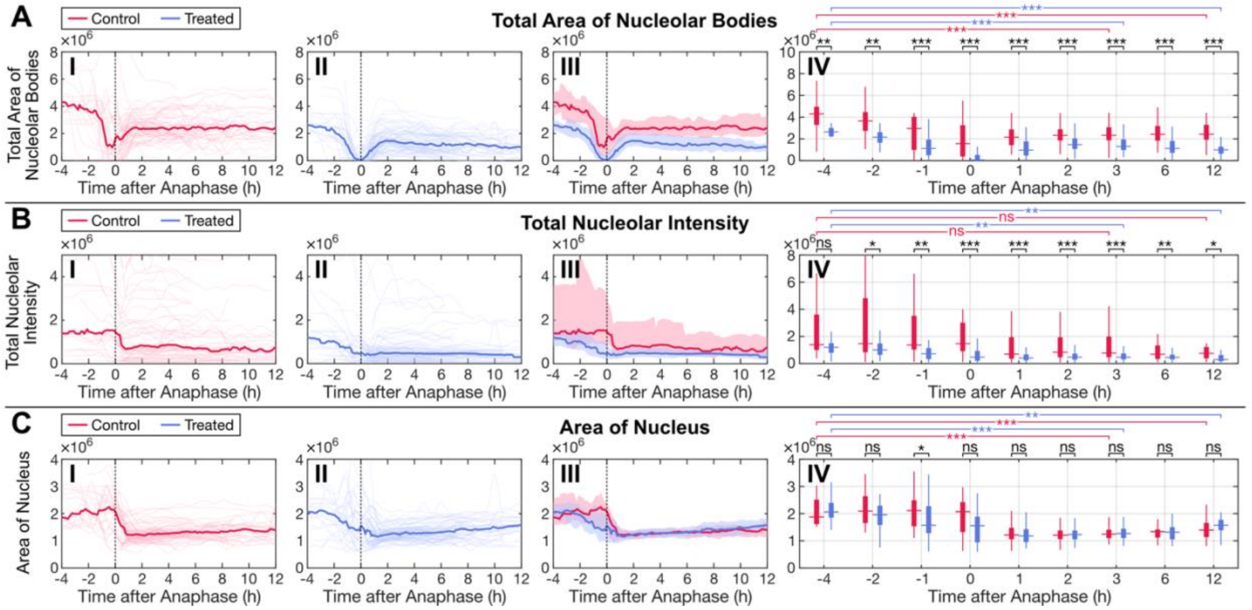


***Supplemental Figure 7 A.*** *Plots of total area of nucleolar bodies,* ***B.*** *total nucleolar intensity, and* ***C.*** *the area of the nucleus, each illustrating* ***I.*** *smoothed raw data for control cells,* ***II.*** *smoothed raw data for ActD treated cells,* ***III.*** *an overlay of the control and treated, and* ***IV.*** *boxplots at different time points.*
